# Deprivation of dietary fiber promotes neuroinflammation and anxiety-like behavior in mice

**DOI:** 10.64898/2026.11.05.764646

**Authors:** Adewale Oyetunde, Dhanya N. Pyaram, Vu Ngo, Chao Li, Hirohito Abo, Jun Zou, Angela M. Mabb, Andrew T. Gewirtz

**Affiliations:** Center for Inflammation, Immunity and Infection, Institute for Biomedical Sciences, Georgia State University, Atlanta, GA, USA; Neuroscience Institute, Center for Diagnostics and Therapeutics, Georgia State University, Atlanta, GA, USA

**Keywords:** dietary fiber, inulin, gut-brain axis, neuroinflammation, microglia, astrocytes, glial bioenergetics, immunometabolism

## Abstract

**Background:** Intestinal health is an important determinant of brain function in that an array of gut perturbants, especially those targeting microbiota, influence behavior. Fermentable dietary fiber nourishes gut microbiota and, consequently, promotes gut health. We hypothesized that depriving mice of fermentable dietary fiber would impact their brain and behavior.

**Methods:** Male C57BL/6J mice were fed standard grain-based chow (GBC), a compositionally defined diet (CDD) which mimics macronutrient content of GBC but lacks fermentable fiber, or CDD supplemented with the fermentable fiber inulin (CDD+INU). Mice were subjected to behavioral testing 6-8 weeks later followed by analysis of the gut brain axis.

**Results:** Relative to GBC-fed mice, CDD-fed mice exhibited low-grade gut inflammation and a reduction in open-field center exploration, signaling heightened anxiety-like behavior. CDD+INU fully reversed these abnormalities arguing they were a consequence of fiber deprivation. Analysis of the gut microbiome and serum metabolome showed that CDD+INU increased oxidative phosphorylation and immune suppressive metabolites. Flow cytometric and transcriptomic analyses of the brain revealed an array of pro-inflammatory alterations in myeloid cells, CD4^+^ T-cells, microglia and astrocytes in mice fed CDD but not CDD+INU. Most notably, microglia and astrocytes from mice fed CDD, but not CDD+INU, displayed reduced mitochondrial respiration and greater glycolytic activity. CDD, but not CDD+INU consumption also reduced the abundance of oligodendrocytes and neurons, and increased hippocampal pro-inflammatory gene expression.

**Conclusions:** Deprivation of fermentable fiber results in coordinated changes in brain immune composition and glial metabolic state that is associated with increased anxiety-like behavior.

## Background

Efforts to better understand determinants of mental health, particularly the post-industrialization increase in a broad array of neuropsychiatric disorders, have led to a growing appreciation of the role of the gut-brain axis, and the gut microbiota therein, in influencing these conditions. For example, exposing mice to high-fat western style diets results in altered gut microbiota composition, elevated pro-inflammatory signaling in gut and/or brain, and alterations in behavior [1]. Our previous work extended this concept to another class of gut microbiota perturbants, namely dietary emulsifiers, which, when fed to mice, altered these parameters in a similar manner [2]. Yet, in our extensive studies that examined the impacts of diet on microbiota and intestinal phenotype, the discrete manipulation that most starkly promoted gut inflammation was deprivation of fermentable dietary fiber [3,4]. Accordingly, we hypothesized that dietary fiber deprivation might also impact brain and/or behavior.

Investigating dietary fiber in animal models is logistically complicated in that grain-based chow (GBC) diet, such as LabDiet 5001, formerly Purina 5001, which have long been used to maintain rodents in research vivaria, are composed of complex, relatively unrefined, and highly variable ingredients, thus precluding altering fiber in a tractable manner [5,6]. Individual batches of the same GBC can range from 12-25% fiber with further variance in the proportion of the fiber that is soluble, and thus readily fermented by gut bacteria, or insoluble, which is resistant to fermentation but yet can have an array of impacts on gut microbiota and, consequently, the intestine. In contrast, compositionally defined diets (CDD) are made from highly refined, quality-controlled ingredients, enabling precise manipulations of nutrients and fiber. CDD that match the average macronutrient content of GBC are widely used as control diets in the nutrition field and are typically quite low in fiber, containing 5% cellulose by weight. This diet can thus be referred to as a low-fiber diet. However, given that CDD is comprised of wholly different ingredients from GBC, attributing impacts of low-fiber CDD to fiber deprivation require reversing such impacts via enriching CDD with fiber. Thus, our conclusion that the intestinal phenotypes that resulted in CDD-fed mice reflected fiber deprivation was based not only on the fact that CDD was low in fiber and induced low-grade inflammation, but also that this phenotype was corrected, in a microbiota-dependent manner, by enriching CDD with the soluble/fermentable fiber inulin [3].

The purpose of the current study was to examine the extent to which the stark impacts of low-fiber CDD and inulin might extend to the central nervous system and, consequently, impact behavior. Indeed, microbial signals are increasingly appreciated to influence microglial maturation and function, and microbiota-derived acetate contributes to microglial metabolic fitness [7,8]. Shi et al. reported that a low-fiber CDD altered the gut microbiota, reduced short-chain fatty acids, increased hippocampal neuroinflammation and microglial synaptic engulfment, and impaired cognition in mice [9], suggesting that dietary content influences fiber neuroimmune pathways. Yet the extent to which such impacts were a consequence of fiber deprivation or another consequence of CDD awaited further study. Hence, we broached this question utilizing GBC, low-fiber CDD and CDD-enriched with inulin. We integrated intestinal phenotyping, microbial and serum metabolomic profiling, brain immunophenotyping, as well as transcriptional, metabolic, and behavioral analyses. We observed that low-fiber diet resulted in a broad array of indices of low-grade neuroinflammation, which was associated with increased anxiety-like behavior, all of which was prevented by inulin arguing these altered phenotypes were indeed a consequence of fiber deprivation.

## Methods

### Animals and dietary intervention

Male C57BL/6J wild-type mice were obtained at 6 weeks of age from The Jackson Laboratory (Bar Harbor, ME, USA). Mice were group-housed at approximately 23 °C under a 12-h light/dark cycle with ad libitum access to food and water and were acclimated for one week before dietary intervention. Animals were assigned to one of three dietary conditions: grain-based chow (GBC), a compositionally defined diet containing cellulose as the principal fiber source (CDD), or the same compositionally defined diet supplemented with inulin (CDD+INU). Diets were maintained for 9 weeks. Behavioral testing began after 6 weeks of dietary exposure. Diet composition, manufacturer and catalog information are provided in Table 1. All procedures were performed at Georgia State University under an Institutional Animal Care and Use Committee-approved protocol.

**Table 1.**
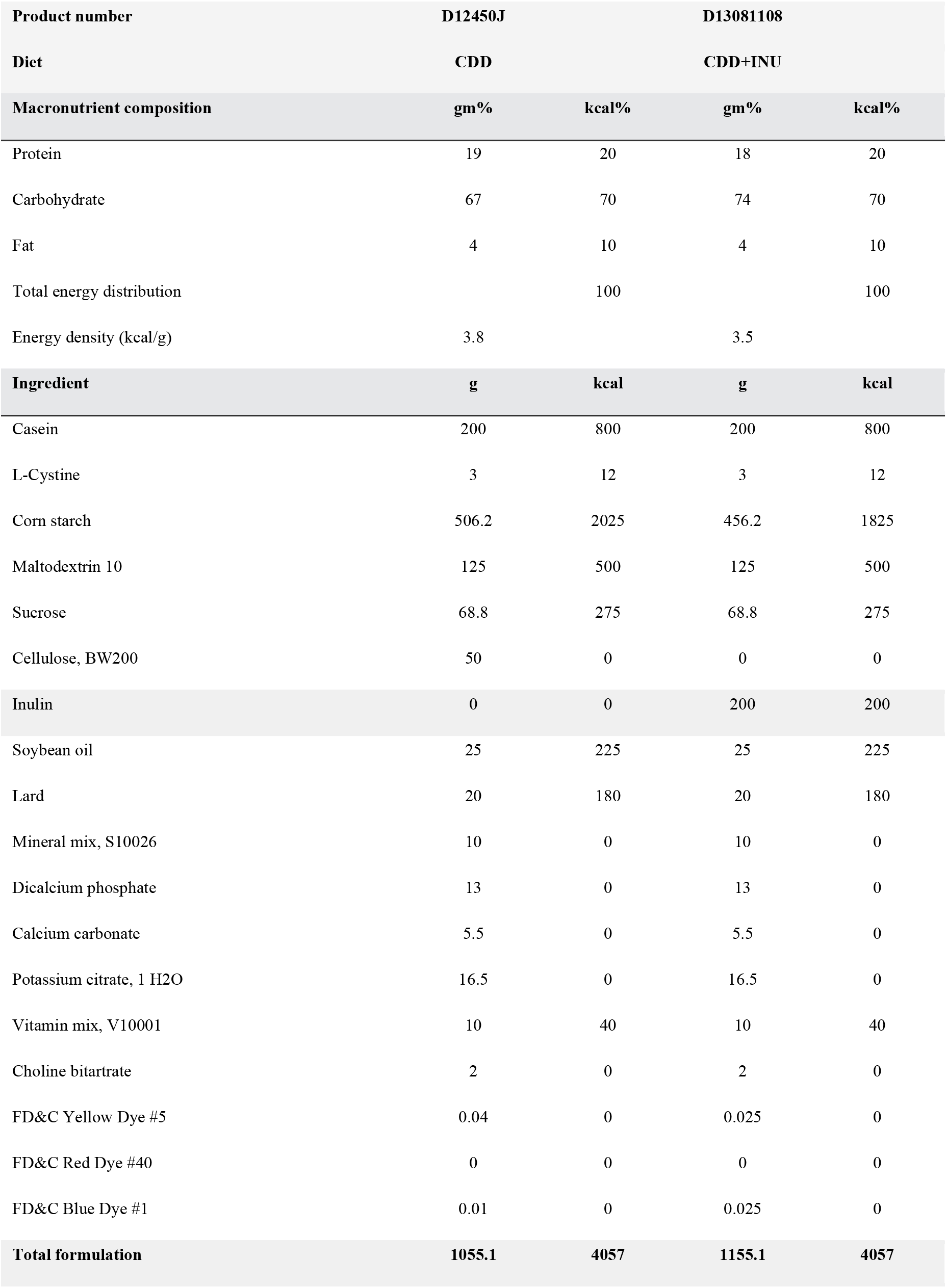
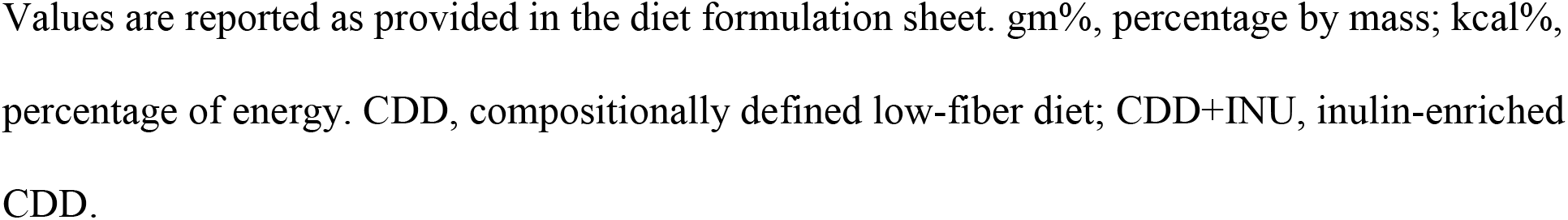
Composition of the experimental low-fiber and inulin-enriched diets.

| **Product number** | **D12450J** |  | **D13081108** |  |
|---|---|---|---|---|
| **Diet** | **CDD** |  | **CDD+INU** |  |
| **Macronutrient composition** | **gm%** | **kcal%** | **gm%** | **kcal%** |
| Protein | 19 | 20 | 18 | 20 |
| Carbohydrate | 67 | 70 | 74 | 70 |
| Fat | 4 | 10 | 4 | 10 |
| Total energy distribution |  | 100 |  | 100 |
| Energy density (kcal/g) | 3.8 |  | 3.5 |  |
| **Ingredient** | **g** | **kcal** | **g** | **kcal** |
| Casein | 200 | 800 | 200 | 800 |
| L-Cystine | 3 | 12 | 3 | 12 |
| Corn starch | 506.2 | 2025 | 456.2 | 1825 |
| Maltodextrin 10 | 125 | 500 | 125 | 500 |
| Sucrose | 68.8 | 275 | 68.8 | 275 |
| Cellulose, BW200 | 50 | 0 | 0 | 0 |
| Inulin | 0 | 0 | 200 | 200 |
| Soybean oil | 25 | 225 | 25 | 225 |
| Lard | 20 | 180 | 20 | 180 |
| Mineral mix, S10026 | 10 | 0 | 10 | 0 |
| Dicalcium phosphate | 13 | 0 | 13 | 0 |
| Calcium carbonate | 5.5 | 0 | 5.5 | 0 |
| Potassium citrate, 1 H2O | 16.5 | 0 | 16.5 | 0 |
| Vitamin mix, V10001 | 10 | 40 | 10 | 40 |
| Choline bitartrate | 2 | 0 | 2 | 0 |
| FD&C Yellow Dye #5 | 0.04 | 0 | 0.025 | 0 |
| FD&C Red Dye #40 | 0 | 0 | 0 | 0 |
| FD&C Blue Dye #1 | 0.01 | 0 | 0.025 | 0 |
| **Total formulation** | **1055.1** | **4057** | **1155.1** | **4057** |

Values are reported as provided in the diet formulation sheet. gm%, percentage by mass; kcal%, percentage of energy. CDD, compositionally defined low-fiber diet; CDD+INU, inulin-enriched CDD.

**Table 2.**
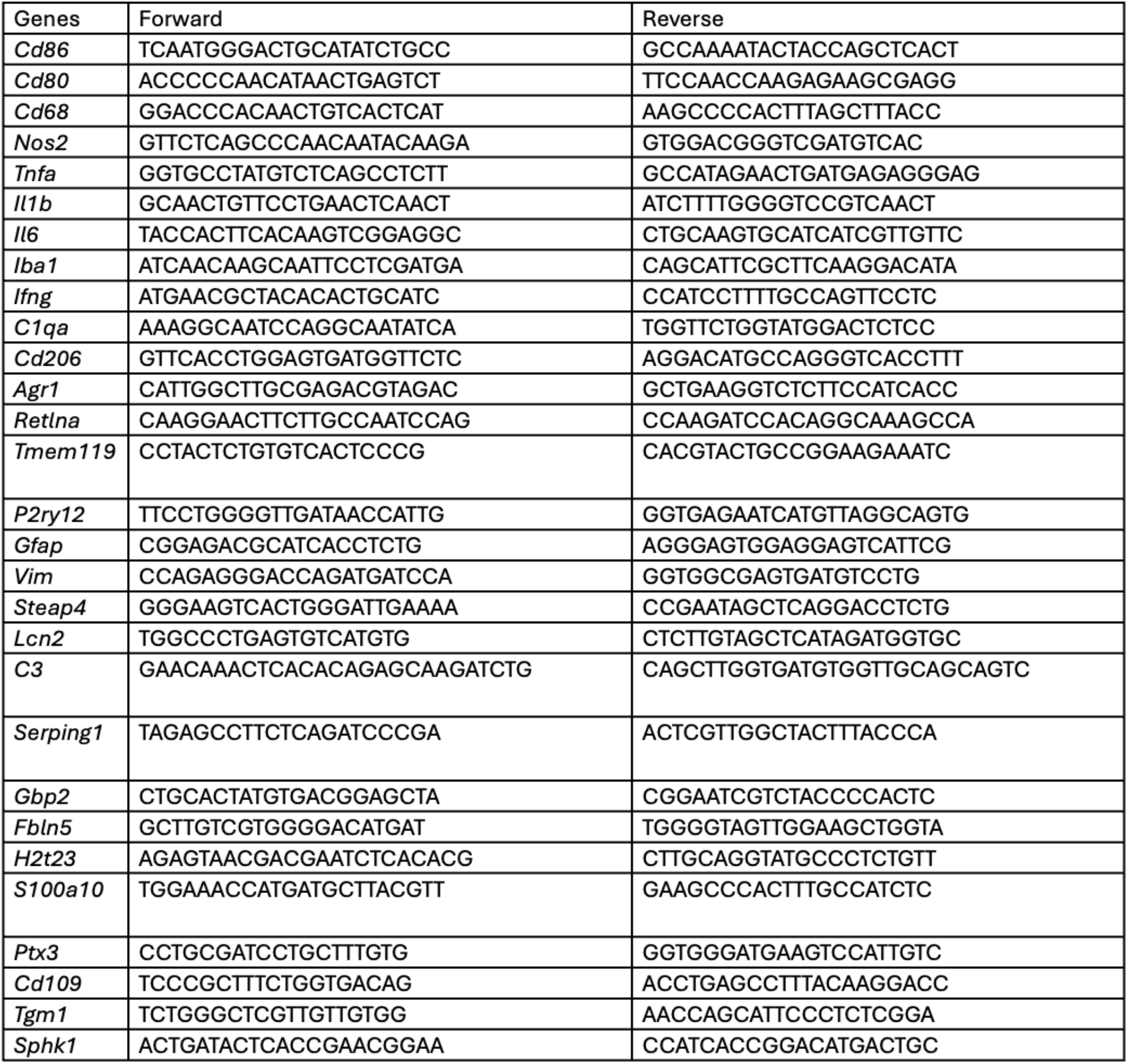
Oligonucleotide.

### Behavioral testing

Open-field testing was performed as previously described [13]. Briefly, mice were acclimated to the testing room for at least 30 min before being placed individually into the open-field arena. Total distance traveled and ambulatory activity were used as locomotor measures, whereas time and/or entries into the predefined center zone were used as indices of center exploration and anxiety-like behavior [13,14]. Forced-swim and tail-suspension testing were performed as described [15,16]. Immobility time during each test period was quantified using ANY-maze automated video-tracking software (Stoelting Co., Wood Dale, IL, USA).

### Tissue collection

At the end of the 9-week feeding period, animals were anesthetized and blood collected before euthanasia. Serum was separated and stored at -80 °C until analysis. Brain, colon and cecum were harvested, and colon length was measured from the cecocolic junction to the anal verge. Cecal contents were collected and frozen. Hippocampi were dissected on ice and snap-frozen for transcript analysis. Animals used for flow cytometry and hippocampal transcriptional analyses were not transcardially perfused; consequently, measurements from these preparations may include residual intravascular cells.

### Brain tissue digestion

Brain tissues were mechanically minced into small fragments and enzymatically digested in RPMI 1640 medium containing collagenase IV (1 mg/mL) and DNase I (50 µg/mL) and 10% fetal bovine serum for 30 min at 37°C with gentle agitation at 200rpm for 20 minutes. Digestion was terminated by adding ice-cold RPMI 1640 supplemented with 10 FBS. The resulting tissue suspensions were passed through 70-µm cell strainers, and centrifuged at 2000rpm for 5 min at 4°C. To remove myelin debris and enrich mononuclear cells, cell pellets were resuspended in 70% isotonic Percoll, overlaid with 37% isotonic Percoll, and centrifuged at 2000rpm for 20 min at room temperature without braking. Cells were collected from the 37%/70% Percoll interface, washed twice with PBS containing 2% FBS, and resuspended in FACS buffer.

### Flow cytometry analysis and isolation of microglia and astrocytes

Isolated cells were incubated with a fixable viability dye to exclude dead cells, followed by Fc receptor blockade with anti-mouse CD16/CD32 antibodies for 15 min at 4°C. Cells were subsequently stained with fluorochrome-conjugated antibodies against the indicated surface markers: CD45 BV605, MHCII BV660, CD11b APC-Cy7, CD11c BV786, O4 PE, NeuN CL594, F4/80 PE, TMEM119 PE-Cy7, ASCA2 APC for 30 min at 4°C in the dark. Following staining, cells were washed twice and resuspended in FACS buffer for flow cytometric analysis. Intracellular staining was done using a FoxP3 staining buffer (eBioscience). Intracellular staining for FoxP3, IL-17A, and IFN-g was performed after re-stimulation of cells with 50 ng/mL phorbol-12-myristate 13 acetate (Sigma) and 750 ng/mL ionomycin (Invitrogen) in the presence of 5 µg/mL Brefeldin A (eBioscience) for 3 hours. Re-stimulated cells were fixed and permeabilized using Fix/Perm buffer set (eBioscience) and stained with antibody IL-17A PE, FoxP3 APC, and IFN-g BV786. Multi-parameter analysis was performed on a CytoFlex (Beckman Coulter) and analyzed with FlowJo software (Tree Star). For sorting of microglia and astrocytes: microglia were defined as CD45^int^CD11b^+^F4/80^++^ and astrocytes were defined as: CD45^-^CD11b^-^O4^-^NeuN^-^CD31^-^ACSA2^+^.

### Metabolic assay of microglia and astrocytes

The metabolic activity of microglia and astrocytes was assessed using the Seahorse XF Cell Mito Stress Test Kit or Seahorse XF Glycolysis Stress Test Kit (Agilent Technologies), according to the manufacturer’s instructions. Microglia and astrocytes were isolated by FACS, seeded into Seahorse cell culture plates at a density of 1 × 10⁴ cells per well, and cultured overnight in complete RPMI 1640 medium. The following day, cells were washed and incubated in Seahorse assay medium (Agilent Technologies) for 1 h before analysis. For the glycolysis stress test, the assay medium was supplemented with 2 mM glutamine, whereas for the mitochondrial stress test, the medium was supplemented with 10 mM glucose and 1 mM pyruvate. All assay reagents were prepared in Seahorse assay medium, with final concentrations of 1.5 μM oligomycin, 4 μM FCCP, and 0.5 μM rotenone/antimycin A for the mitochondrial stress test, and 10 mM glucose, 1 μM oligomycin, and 50 mM 2-deoxy-D-glucose (2-DG) for the glycolysis stress test. Oxygen consumption rate (OCR) and extracellular acidification rate (ECAR) were measured using a Seahorse XF analyzer.

### Quantitative real-time PCR

RNA from FACS sorted microglia cells (hemibrain) and FACS sorted astrocytes cells (hemibrain) were isolated using the Qiagen RNeasy Mini Kit with on-column DNase digestion according to manufacturer’s protocol. cDNA was generated using the Superscript First Strand Synthesis kit for RT-PCR (Invitrogen). RT-qPCR was performed with SYBR Green using StepOnePlus PCR system (Applied Biosystem), and gene expression was normalized to GAPDH. RNA was isolated from hippocampal tissue and analyzed by one-step quantitative RT-PCR. Expression of *Il6, Tnf, Il1b, Ifng, Cd68, Aif1*, was quantified and normalized to *Gapdh*. Primers listed in Table S2.

16S rRNA gene sequencing and microbial community analysis

Fecal bacterial communities were analyzed by 16S rRNA gene sequencing as previously described [17]. Briefly, DNA was extracted from frozen fecal samples from CDD and CDD+INU mice (n = 4/group) using the PowerSoil-htp kit (MO BIO Laboratories, Carlsbad, CA, USA) with mechanical disruption by bead beating. The V4 region of the bacterial 16S rRNA gene was amplified using the conserved bacterial primers 515F (5′-GTGCCAGCMGCCGCGGTAA-3′) and 806R (5′-GGACTACHVGGGTWTCTAAT-3′). Amplicons were sequenced on an Illumina MiSeq platform using paired-end 2 × 250-bp sequencing. Sequence data were processed using QIIME 2, with DADA2 used for quality filtering, denoising, merging, and chimera removal. Taxonomy was assigned using the SILVA 138 reference database.

### Prediction of microbial metagenomic functions using PICRUSt2

The functional potential of the intestinal microbiota from mice fed with GBC, CDD and CDD+INU was predicted from 16S rRNA sequencing data using PICRUSt2 through the Galaxy Europe platform (https://usegalaxy.eu). Briefly, representative amplicon sequence variant (ASV) sequences and their corresponding abundance tables were uploaded in FASTA and BIOM formats. PICRUSt2 was used to predict microbial gene content based on the phylogenetic relationships between ASVs and reference genomes. Predicted gene abundances were normalized by estimated 16S rRNA gene copy numbers and annotated using Enzyme Commission (EC) numbers and KEGG Orthologs (KOs).

### Untargeted serum metabolomics

Untargeted GC-MS metabolomic profiling was performed in CDD and CDD+INU mice (n = 5/group). GBC animals were not included in this analysis. Serum metabolites were extracted, derivatized and analyzed by GC-MS. Peak abundances were normalized to the internal standard and sample volume. Metabolites were identified by retention-index and spectral matching against [library TO CONFIRM]. Data were log-transformed and scaled before principal-component and hierarchical-clustering analyses. False-discovery-rate correction was applied to individual metabolite comparisons.

### Short-chain fatty acid analysis

Levels of acetate, propionate, and butyrate in cecal contents collected from non-fasted mice were measured by gas chromatography as previously described [18].

### Statistical analysis

Data are presented as mean ± SEM unless otherwise indicated. Three-group comparisons were analyzed using one-way ANOVA followed by Tukey’s multiple-comparisons test where assumptions for parametric testing were satisfied. Two-group analyses used appropriate two-sided comparisons. Repeated body-weight measurements were analyzed using repeated-measures analysis. False-discovery-rate correction was applied to untargeted metabolomic analyses. Microbiome analyses were considered exploratory because of the small sequencing cohort (n = 4/group). Exact biological sample sizes are reported with each final figure. Statistical significance was defined as p < 0.05.

## Results

### Deprivation of dietary fiber promotes anxiety-like behavior

We first assessed consequences of feeding mice standard grain-based chow (GBC), which is naturally rich in fiber (12-23% by weight) versus a highly refined compositionally-defined diet (CDD), comprised of only 5% fiber, all insoluble (cellulose), which is widely used as a control diet in nutrition research. Groups of 6-week-old male C57BL/6 mice (n=10) remained on GBC diet or were switched to CDD for a 9-week period during which body weight was monitored and they were subjected to behavioral testing **(Fig 1A)**. Specifically, mice were subjected to the open field testing (at 6 weeks) to assess anxiety-like behavior, followed by tail suspension and forced swim testing (7 and 8 weeks, respectively) as a means of probing for depressive-like behaviors. Mice were then euthanized at 9 weeks for downstream gut-brain axis analyses. In accord with previous studies, CDD resulted in mild weight gain **(Fig 1B)**, which herein was transient, potentially reflecting the longer duration of the current study relative to our previous work or that the behavioral testing causes more stress in CDD-fed mice. In any event, a cardinal feature of CDD-feeding, namely colon shortening **(Fig 1C)**, which reflects gut inflammation, was evident in euthanized mice indicating the previously observed consequences of CDD on the intestine were maintained throughout the experiment. Open field testing **(Fig 1D)** found that CDD-fed mice displayed significantly greater distance traveled and reduced time spent in the center (p = 0.03 and 0.01 by t-test) while tail suspension and forced swim testing revealed a trend that was not statistically significant between GBC- and CDD-fed mice **(Fig 1E,F)**. These results suggested that CDD feeding had promoted anxiety-like behavior but had 2 key limitations, namely that applying a Bonferroni correction to our results would have made the differences only moderately significant and, furthermore, our inability to discern the extent to which impacts of CDD on behavior were a consequence of its lack of fiber.

**Figure 1.**
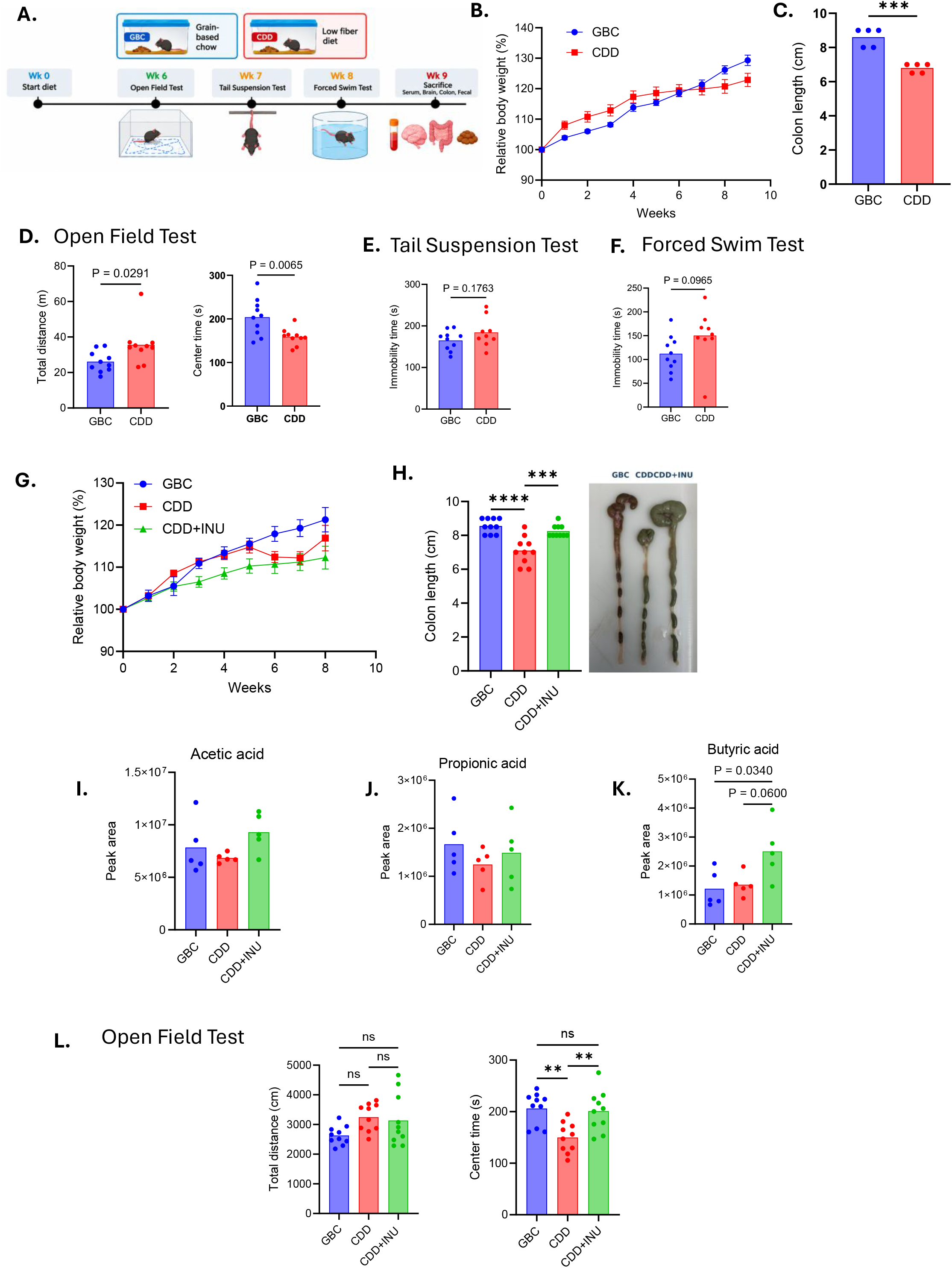
Inulin supplementation attenuates physiological and behavioral alterations associated with consumption of a compositionally defined low-fiber diet. (A) Experimental timeline showing dietary interventions, behavioral testing, and sample collection. (B–F) Comparison of grain-based chow (GBC)- and compositionally defined low-fiber diet (CDD)-fed mice: (B) relative body weight, (C) colon length, (D) open-field total distance and center time, (E) tail suspension test immobility, and (F) forced swim test immobility. (G–L) Assessment of GBC-, CDD-, and inulin-supplemented CDD (CDD+INU)-fed mice: (G) relative body weight, (H) colon length and representative gastrointestinal morphology, (I–K) serum acetate, propionate, and butyrate peak areas, respectively, and (L) open-field total distance and center time. Data are presented as group means with individual biological replicates where applicable. Statistical significance: P < 0.05, P < 0.01, P < 0.001, P < 0.0001; ns, not significant.

To surmount these limitations, we repeated the experiment with a new cohort of mice and included a third group receiving CDD enriched with the fermentable fiber inulin (CDD+INU), while focusing a priori on the prediction that CDD, but not CDD+INU, would promote anxiety-like behavior. All three dietary groups exhibited progressive increases in body weight over the 8-week intervention, with CDD+INU-fed mice showing the lowest relative weight gain, followed by CDD- and GBC-fed mice **(Fig 1G)**. In accord with previous studies, CDD+INU feeding prevented the stark morphological changes, including colon-shortening induced by CDD **(Fig 1H)**. Moreover, such protection conferred by enriched CDD with inulin was associated with increased levels of cecal short-chain fatty acids (SCFA), particularly butyrate **(Fig 1I-1K)**. Most importantly, an identical pattern of differences was observed in open field testing, specifically increased total distance traveled and reduced center time in mice fed CDD but not CDD+INU **(Fig 1L)**. These results indicated that, as a result of its lack of fermentable fiber, CDD promoted anxiety-like behavior in mice.

### Untargeted omics analysis suggests broad changes in microbiota-related brain-relevant metabolites

We hypothesized that the impact of dietary fiber on behavior reflected a role for microbiota-regulated metabolites. This group of metabolites is increasingly appreciated to extend well beyond SCFA, prompting us to use an untargeted approach. We initially considering analyzing metabolomes from mice fed GBC, CDD, and CDD+INU but found that the differences between GBC and CDD were vast, reflecting the wholly different ingredients that comprise these diets. Hence, we focused on identifying metabolites that differed in abundance due to differences in fermentable fiber content by performing untargeted metabolomic analysis of serum from mice fed CDD or CDD+INU **(Fig 2A)**. Overall assessment of the data via principal coordinate analysis showed significant clustering (PERMANOVA=0.032) of serum metabolomes based on whether mice consumed CDD versus CDD+INU indicating addition of fermentable fiber to a low-fiber diet had a significant impact on overall metabolome **(Fig 2B)**. Such overall difference was driven by approximately equal numbers of metabolites being increased and decreased in abundance **(Fig 2C)** although, there was considerable mouse to mouse variation and thus, not surprisingly, the large number of metabolites detected by the untargeted assay rendered all observed differences in individual metabolites non-significant following Bonferroni correction. Nonetheless, a number of differences in metabolites arbitrarily selected based on uncorrected statistical significance and/or previous reports that they were mediators of the gut brain axis were noted including urea, creatine, glutamic acid, glycolytic/TCA-related intermediates, β-hydroxybutyrate, oleic acid, and linoleic acid **(Fig 2D-O)**. With the exception of β-hydroxybutyrate, which is reported to lower anxiety-like behavior [10], but yet was higher in CDD+INU fed mice, results were consistent with the hypothesis that fermentable fiber drives microbiota-mediated changes in the metabolome that reduce anxiety-like behavior.

**Figure 2.**
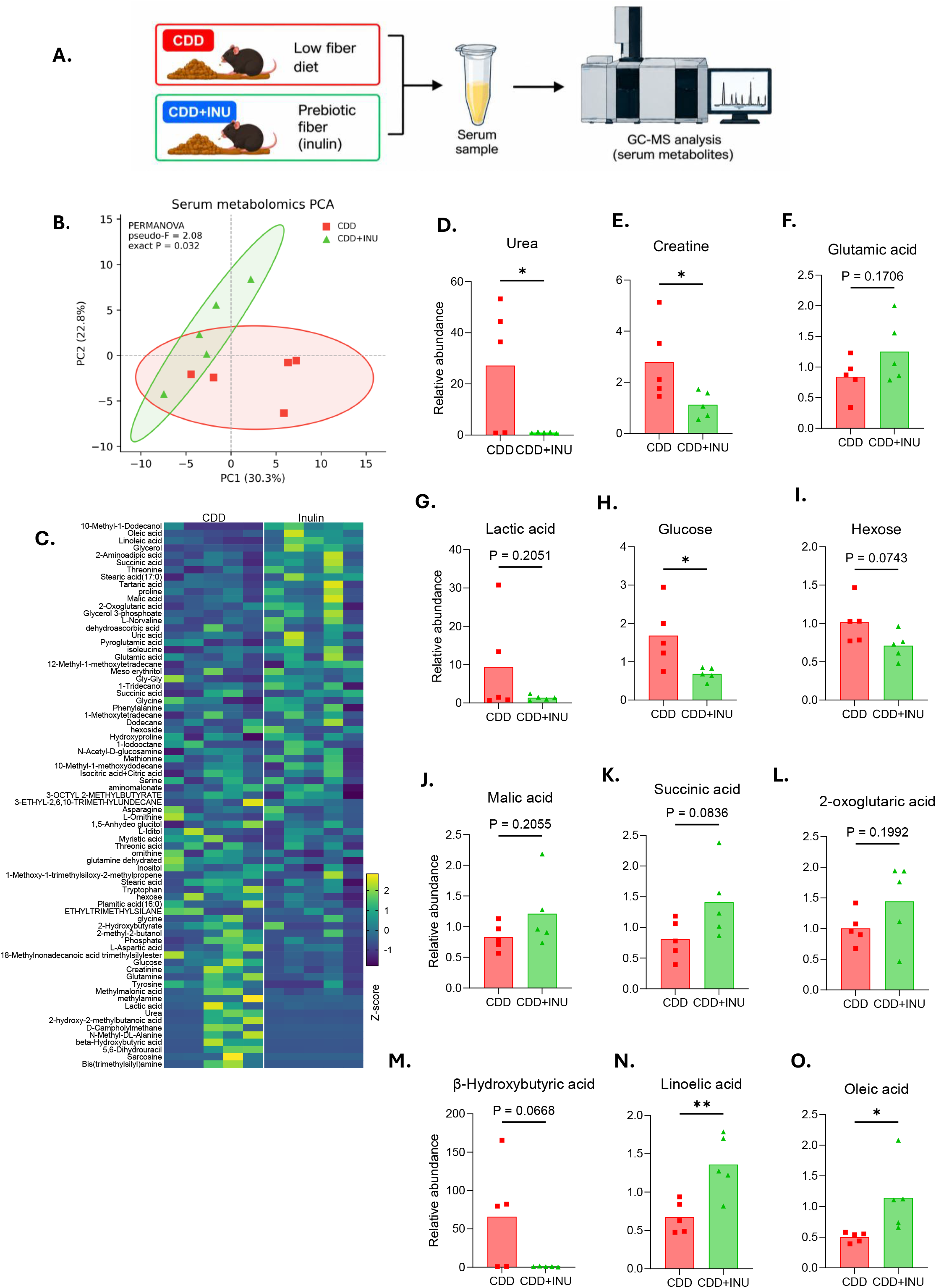
Inulin supplementation modulates serum metabolomic profiles in mice fed a compositionally defined low-fiber diet. Serum metabolites were analyzed by gas chromatography–mass spectrometry (GC–MS) in CDD- and CDD+INU-fed mice. (A) Schematic of serum collection and GC–MS analysis. (B) Principal component analysis (PCA) of serum metabolomic profiles (PERMANOVA, pseudo-F = 2.08, P = 0.032). (C) Heatmap showing relative metabolite abundance across individual samples, expressed as Z-scores. (D–O) Relative abundance of selected metabolites: (D) urea, (E) creatine, (F) glutamic acid, (G) lactic acid, (H) glucose, (I) hexose, (J) malic acid, (K) succinic acid, (L) 2-oxoglutaric acid, (M) β-hydroxybutyric acid, (N) linoleic acid, and (O) oleic acid. Bars represent group means with individual biological replicates. Individual metabolite comparisons were exploratory and did not remain significant after multiple-comparison correction. Statistical significance shown for unadjusted comparisons: P < 0.05, P < 0.01; exact P values are indicated where applicable.

We next probed potential relationships between the serum metabolomes and microbiomes of CDD and CDD+INU mice, analyzing fecal microbiomes by 16S RNA sequencing. We used an n=4 based on earlier studies that had analyzed impacts of these diets on microbiota albeit in studies of moderately shorter duration. Results were largely reminiscent of our earlier findings with mice exhibiting diet-based clustering by both weighted Unifrac and Bray-Curtis analysis **(Fig 3 A,B)** that associated with a reduction in α-diversity (species richness) **(Fig 3C),** and an array of specific taxonomic differences including increased abundance of *Muribaculaceae*, *Roseburia*, and *Lactobacillus hominis* and reduced abundance of *Eggerthellaceae*, *Turicibacter*, and *Erysipelatoclostridiaceae* **(Fig 3D-E).**

**Figure 3.**
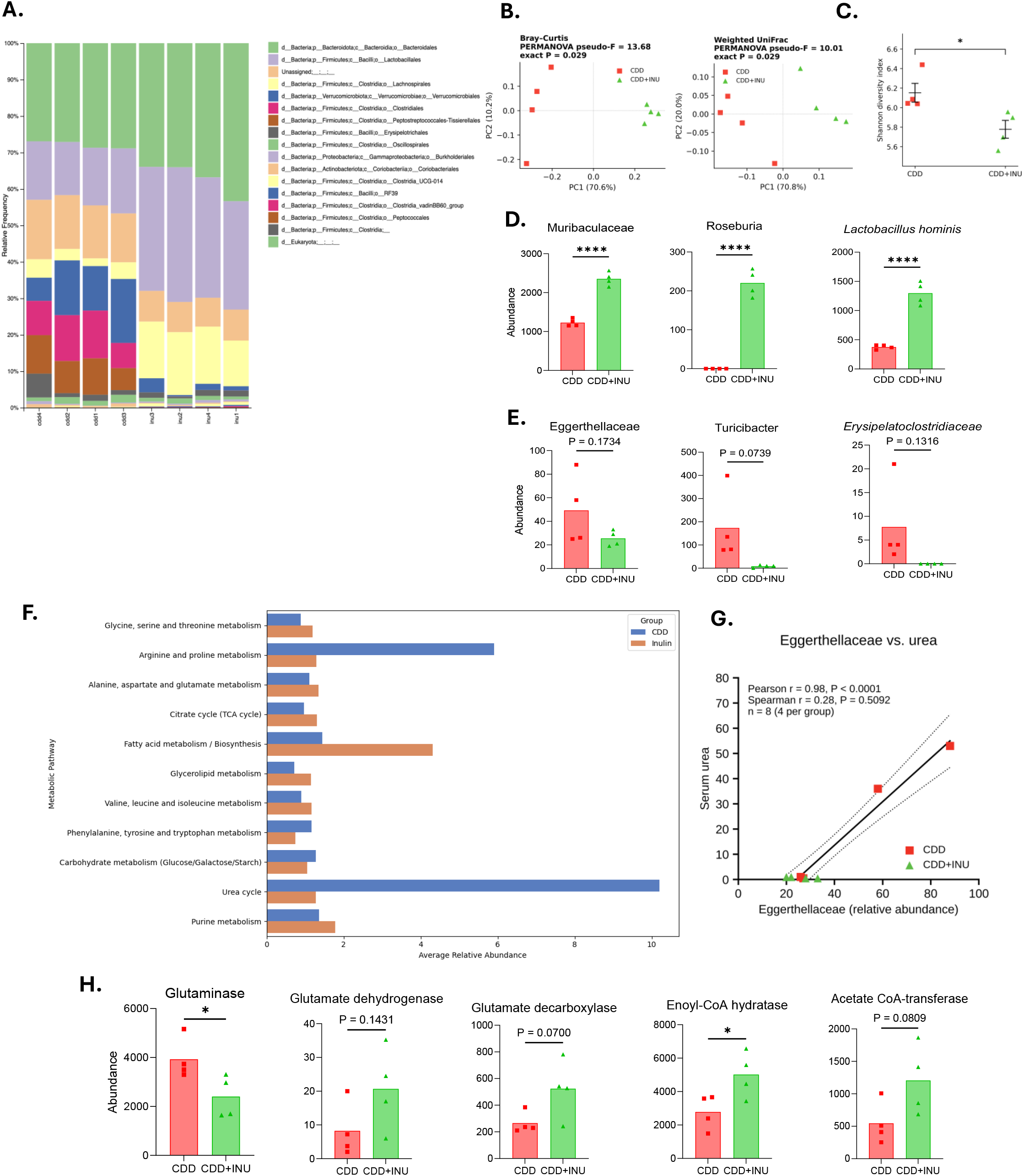
Inulin supplementation is associated with restructuring of the fecal microbiota and its predicted metabolic potential in CDD-fed mice. Fecal microbiota were characterized by 16S rRNA gene sequencing in CDD- and CDD+INU-fed mice. (A) Stacked bar plots showing relative taxonomic abundance across individual samples. (B) Principal coordinate analysis (PCoA) based on Bray–Curtis and weighted UniFrac distances (PERMANOVA: pseudo-F = 13.68, P = 0.029 and pseudo-F = 10.01, P = 0.029, respectively). (C) Shannon alpha diversity. (D–E) Relative abundance of selected bacterial taxa, including *Muribaculaceae, Roseburia, Lactobacillus hominis, Eggerthellaceae, Turicibacter, and Erysipelatoclostridiaceae*. (F) Predicted microbial metabolic pathways. (G) Exploratory association between *Eggerthellaceae* abundance and serum urea (Pearson r = 0.98, P < 0.0001; Spearman r = 0.28, P = 0.5092; n = 8). (H) Predicted microbial enzyme abundances associated with amino acid and energy metabolism. Functional predictions represent inferred metabolic potential rather than directly measured activity. Bars represent group means with individual biological replicates. Statistical significance is indicated in the panels.

We used PICRUSt2 to predict functional consequences of these microbiomes. The most prominent prediction this algorithm yielded was that CDD+INU feeding would starkly reduce flux through the urea cycle, a result which comported with the metabolomics analysis, which found markedly lower levels of urea in mice fed CDD+INU. Hence, we probed the extent to which abundance of any of the above noted taxonomic changes correlated with urea abundance. Small sample size notwithstanding, results suggested that *Eggerthellaceae*, a taxon previously associated with stress-and anxiety-like behavior, met this criterion [11,12]. In light of our metabolomics analysis suggesting differences related to glutamate and TCA cycle intermediated, we also used PICRUSt2 to predict levels of enzymes relevant to these metabolites. It predicted lower glutaminase and higher glutamate dehydrogenase, glutamate decarboxylase, enoyl-CoA hydratase, and acetate CoA-transferase abundance **(Fig 3F-H)**. Collectively, these omics analysis, although exploratory, accorded with the possibility that fermentable fiber levels might impact the abundance of microbiota-related metabolites that are capable of influencing brain and behavior.

### Fiber deprivation alters brain immune cell populations promoting low-grade inflammation

We next examined the consequences of manipulating dietary fiber on the brain. Whole hemibrains were isolated from mice that had been fed GBC, CDD, or CDD+INU over the 9-week period, during which they were subjected to behavioral testing. Such hemibrains were digested to single cell suspensions and assayed by flow cytometry **(Fig 4A)**. We particularly looked for indices of low-grade inflammation previously observed in the gut of CDD-fed mice, hypothesizing that such low-grade inflammation might extend to the brain. Data are shown as representative flow plots, in which individual cell types are quantitated relative to the total leukocytes (CD45**^+^** cells) in the brain and then in bar plots showing absolute abundance in brain. In accord with our hypothesis, relative to mice fed GBC, feeding of CDD, but not CDD+INU, resulted in increased brain abundance of innate immune cells, namely neutrophils **(Fig 4B)** and inflammatory monocytes **(Fig 4C)**, while levels of dendritic cells, which link innate to adaptive immunity and are regulated more at the functional level than by abundance, did not change **(Fig 4D)**. Furthermore, fiber deprivation also impacted levels of T-helper lymphocytes in a manner consistent with inflammation. Specifically, mice fed CDD but not CDD+INU displayed elevated levels of major pro-inflammatory Th subsets, namely Th1 and Th17 cells, and a reduction in Foxp3**^+^** regulatory T-cells **(Fig 4E-G)**. An important caveat to these data is that we cannot be certain of whether the leukocytes we quantitated were truly in the brain parenchyma or the cerebral blood vessels. Nonetheless, these data argue the brain environment of fiber deprived mice exhibited low-grade inflammation.

**Figure 4.**
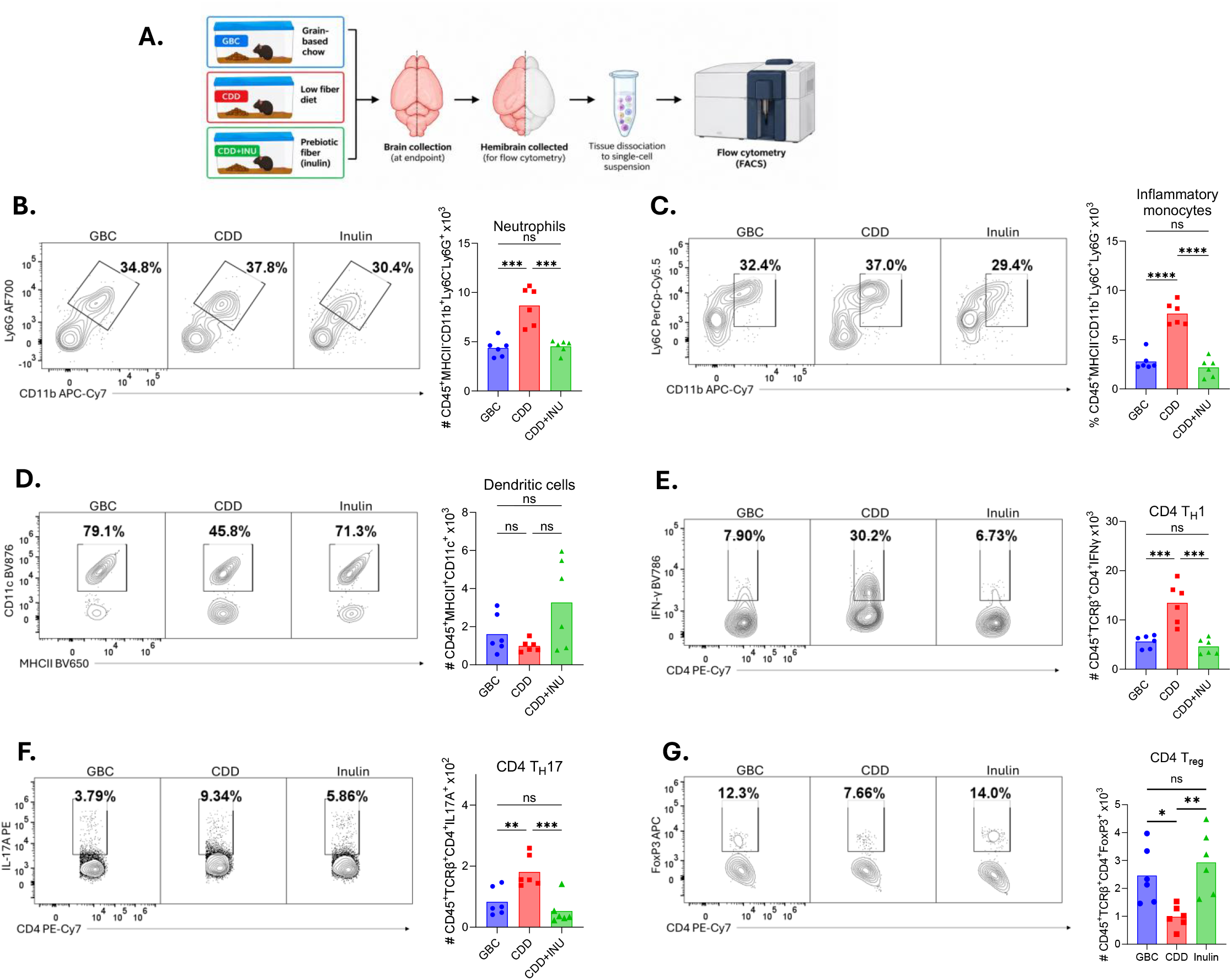
Inulin supplementation attenuates CDD-associated alterations in brain-associated immune-cell populations. Brain-associated immune cells were characterized by flow cytometry in GBC-, CDD-, and CDD+INU-fed mice. (A) Schematic of brain collection, hemibrain processing, and flow cytometric analysis. (B–D) Representative plots and quantification of (B) neutrophils (CD45⁺MHCII⁻CD11b⁺Ly6C⁺Ly6G⁺), (C) inflammatory monocytes (CD45⁺MHCII⁻CD11b⁺Ly6C⁺Ly6G⁻), and (D) dendritic cells (CD45⁺MHCII⁺CD11c⁺). (E–G) Representative plots and quantification of CD4⁺ T-cell subsets: (E) IFN-γ⁺ Th1 cells, (F) IL-17A⁺ Th17 cells, and (G) Foxp3⁺ regulatory T cells. Bars represent group means with individual biological replicates. Brains were not perfused before collection; thus, analyzed populations may include vascular-associated cells. Statistical significance: P < 0.01, P < 0.001, P < 0.0001; ns, not significant.

### Fiber deprivation results in a more pro-inflammatory microglial phenotype

Brain-resident macrophages, referred to as microglial cells are the predominant cell type charged with protecting the brain. Like macrophages in other tissues, microglia exhibit surface expression of CD45, CD11b, and F4/80. Furthermore, most microglial cells also express TMEM119, which indicates they are long-time brain residents. In contrast, microglia lacking expression of this marker are more likely to have recently arrived in the brain and are often to referred as infiltrating macrophages. Assay of microglia by FACS **(Fig 5A)** found that levels of total microglia did not differ based on diet **(Fig 5B)** but feeding CDD but not CDD+INU resulted in a modest reduction of brain-resident TMEM119**^+^** microglia and a corresponding increase in infiltrating macrophages **(Fig 5C).** Furthermore, feeding CDD, but not CDD+INU resulted in an increased proportion of disease-associated microglia expressing an M1-like phenotype and decreased the number of activated-response microglia expressing an M2-like phenotype **(Fig 5D).** These differences are all classic features of low-grade inflammation. Moreover, assay of microglial gene expression by qRT-PCR found increased expression of an array of pro-inflammatory genes associated with an M1 phenotype and decreased expression of anti-inflammatory genes associated with an M2 phenotype in mice fed CDD but not CDD+INU **(Fig 5E)**. Lastly, we performed real-time metabolic analysis (Seahorse) in microglial cells. Relative to mice fed GBC, those fed CDD, but not CDD+INU exhibited increased use of glycolysis **(Fig 5F)**, which is an immune cell hallmark in a more activated state. Thus, consumption of CDD, but not CDD+INU altered the phenotype of microglia towards a more pro-inflammatory state.

**Figure 5.**
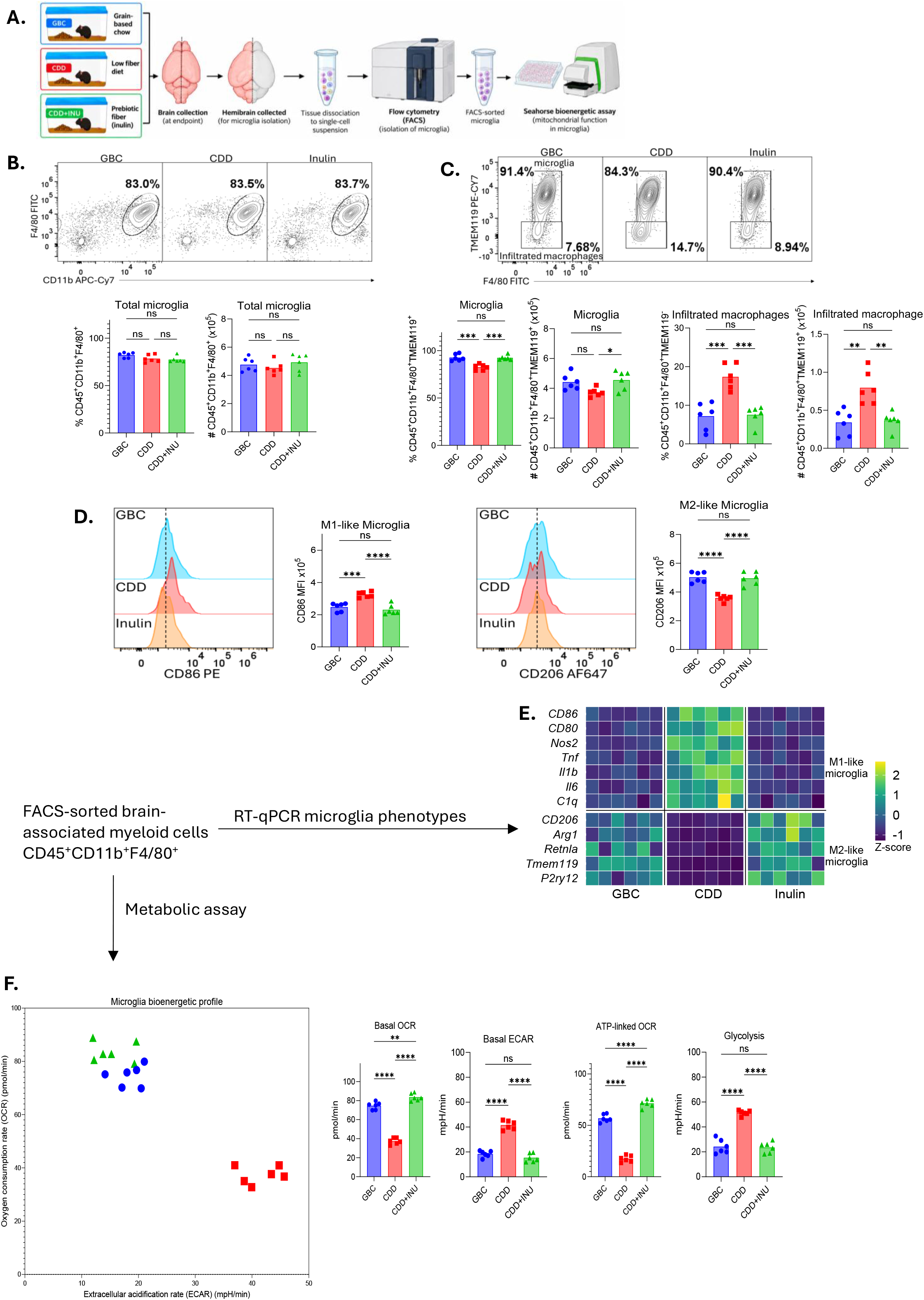
Inulin supplementation attenuates CDD-associated microglial inflammatory activation and metabolic reprogramming. Brain-associated myeloid cells were isolated from mice fed grain-based chow (GBC), a compositionally defined low-fiber diet (CDD), or CDD supplemented with inulin (CDD+INU). (A) Schematic of brain tissue processing, fluorescence-activated cell sorting (FACS), and Seahorse metabolic analysis. (B) Representative flow cytometry plots and quantification of CD45⁺CD11b⁺F4/80⁺ myeloid cells. (C) Frequencies and counts of TMEM119⁺ microglia and TMEM119⁻ myeloid cells. (D) Representative histograms and quantification of CD86 and CD206 mean fluorescence intensity (MFI). (E) Heatmap of inflammatory and anti-inflammatory gene expression in FACS-isolated cells, assessed by RT-qPCR. (F) Seahorse bioenergetic profiling showing oxygen consumption rate (OCR), extracellular acidification rate (ECAR), basal respiration, ATP-linked respiration, and glycolysis. Bars represent group means, with individual data points indicating biological replicates. Statistical significance: P < 0.05, P < 0.01, P < 0.001, P < 0.0001; ns, not significant.

### Astrocytes exhibit parallel transcriptional and bioenergetic remodeling

We next examined impacts of modulating dietary fiber on astrocytes, which were identified as CD45^-^CD11b^-^ACSA2^+^O4^-^cells **(Fig 6A)**. Fewer live astrocytes were recovered in mice fed CDD, but not CDD+INU **(Fig 6B)**, suggesting the possibility that reduced abundance of these cells or their capacity to withstand the isolation procedures, although the statistical significance of the difference was modest. Gene expression of FACS-isolated astrocytes was assayed by qRT-PCR focusing on panels of genes widely used as signatures of pan-reactivity (*Gfap*, *Vim*, *Steap4*, and *Lcn2*), neurotoxicity (*c3*, *Serping1*, *Gbp2*, *Fbln5*, and *H2-T23*), and neuroprotection (*S100a10*, *Ptx3*, *Cd109*, *Tgm1*, and *Sphk1*). Relative to mice maintained on a standard GBC diet, astrocytes from mice fed CDD, but not CDD+INU exhibited more pan-reactive and neurotoxic gene expression patterns and correspondingly reduced expression of genes that mediate neuroprotection **(Fig 6C)**. Concomitantly, astrocytes from CDD-fed mice were more glycolytic whereas astrocytes from mice fed GBC or CDD+INU relied more on oxidative phosphorylation **(Fig 6D)**. Collectively, these results indicate that fiber deprivation compromises the neuroprotective properties of astrocytes.

**Figure 6.**
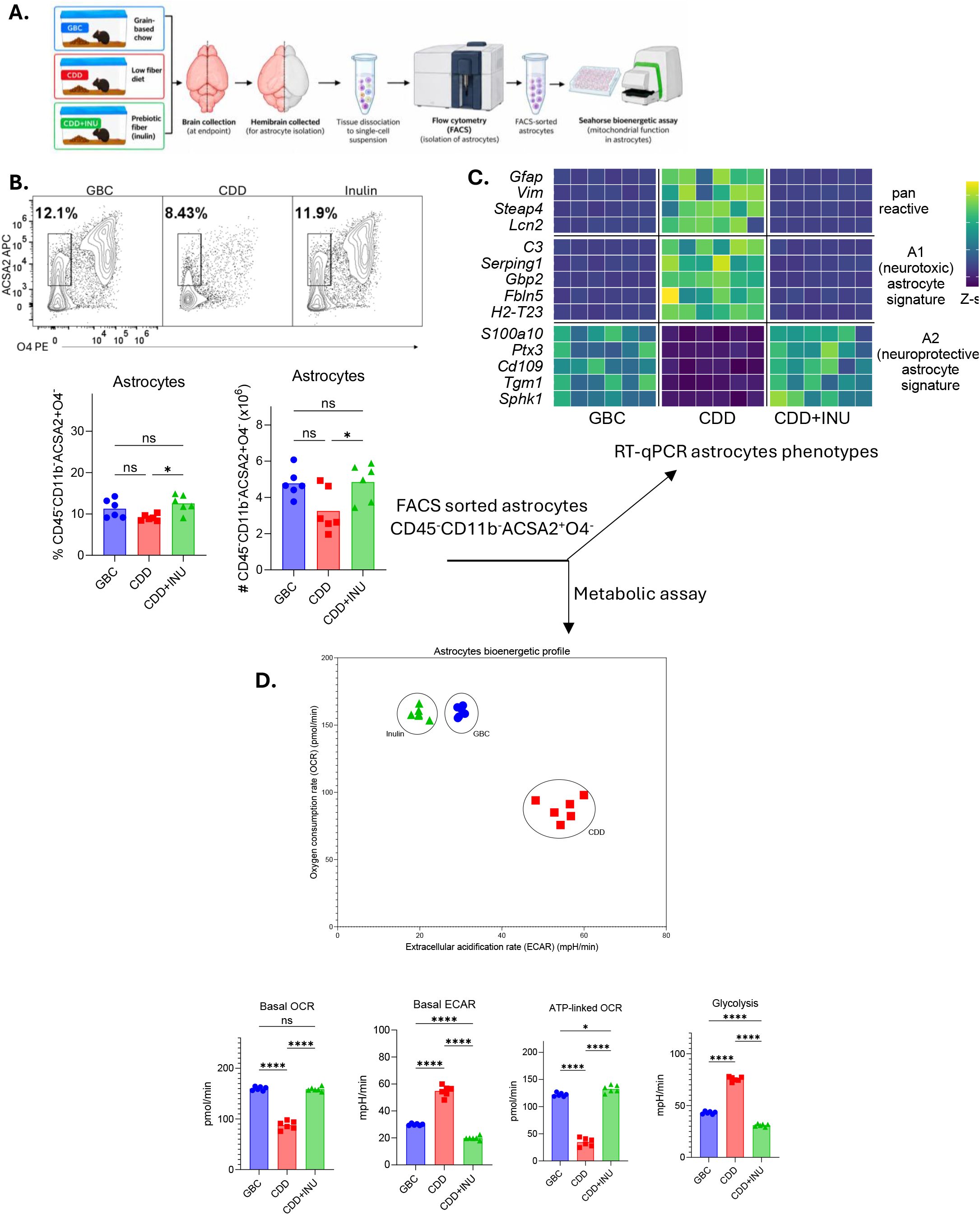
Inulin supplementation attenuates CDD-associated astrocyte inflammatory activation and metabolic reprogramming. Astrocytes were isolated from mice fed grain-based chow (GBC), a compositionally defined low-fiber diet (CDD), or CDD supplemented with inulin (CDD+INU). (A) Schematic of brain tissue processing, fluorescence-activated cell sorting (FACS), and Seahorse metabolic analysis. (B) Representative flow cytometry plots and quantification of CD45⁻CD11b⁻ACSA2⁺O4⁻ astrocytes. (C) Heatmap of pan-reactive, neurotoxicity-associated (A1), and neuroprotection-associated (A2) gene expression signatures assessed by RT-qPCR and expressed as Z-scores. (D) Seahorse bioenergetic profiling showing oxygen consumption rate (OCR), extracellular acidification rate (ECAR), basal OCR, basal ECAR, ATP-linked OCR, and glycolysis. Bars represent group means, with individual points indicating biological replicates. Statistical significance: P < 0.05, P < 0.01, P < 0.0001; ns, not significant.

### Reduced neural cells and increased hippocampal pro-inflammatory gene expression in fiber-deprived mice

We hypothesized that the above-described changes in immune cells and astrocytes might have impeded wellness of the cells they support, namely oligodendrocytes and neurons. To investigate this notion, we quantitated levels of oligodendrocytes (CD45-CD11b-ACSA2-O4+ cells and neurons (CD45-NeuN+ cells) by flow cytometry **(Fig 7A)**. This approach only quantifies live cells and thus cannot distinguish whether any observed differences in cell number existed in the brain or occurred during tissue dissociation and recovery. This caveat notwithstanding, feeding CDD, but not CDD+INU, resulted in reduced abundance of both oligodendrocytes and neurons (Fig 7 B,C).

**Figure 7.**
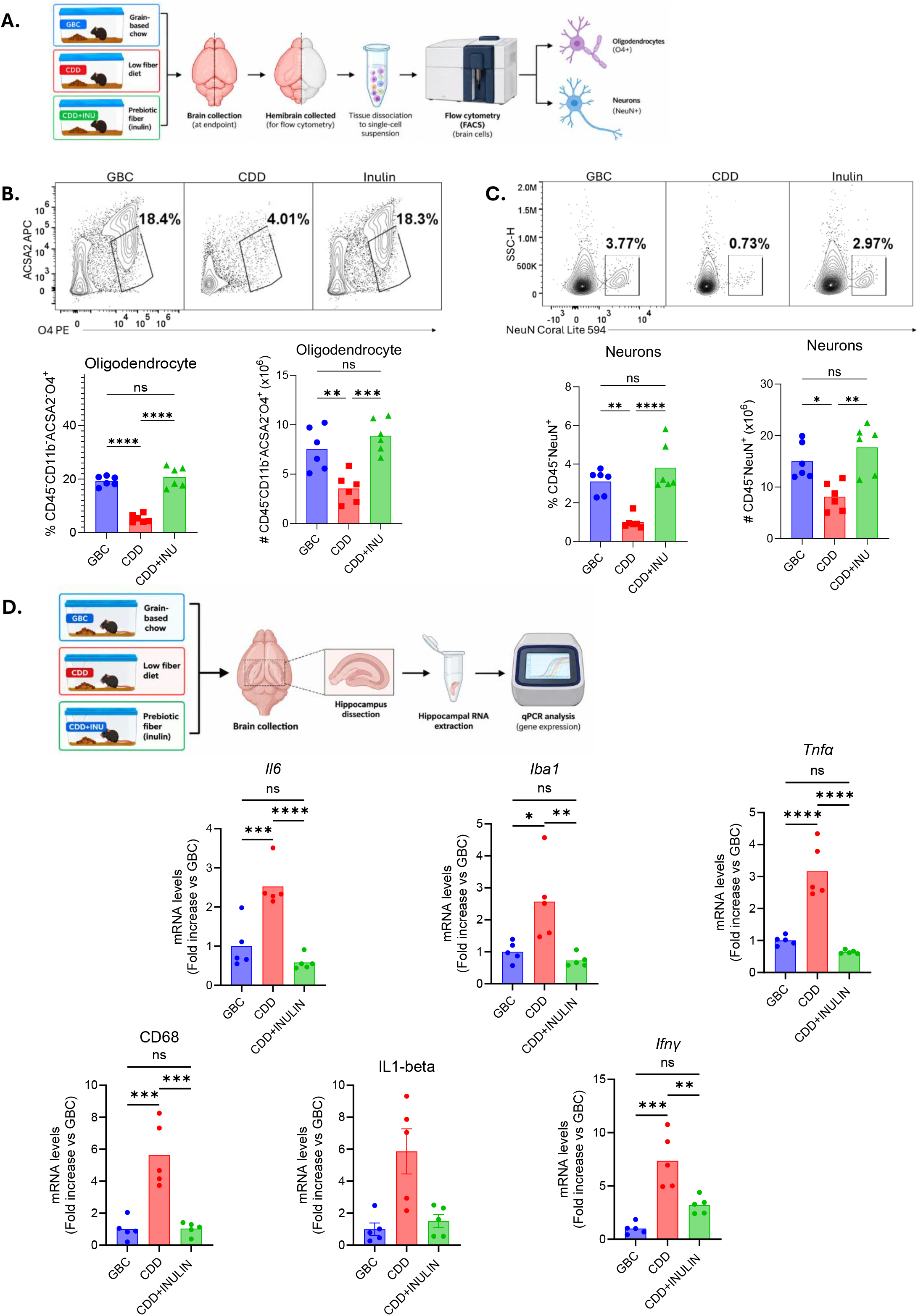
Dietary fiber enrichment attenuates CDD-associated reductions in recoverable oligodendrocytes and neurons and hippocampal inflammatory gene expression. Mice were fed grain-based chow (GBC), a compositionally defined low-fiber diet (CDD), or CDD supplemented with inulin (CDD+INU). (A) Schematic of brain tissue processing and flow cytometric analysis of oligodendrocytes and neurons. (B–C) Representative flow cytometry plots and quantification of (B) O4⁺ oligodendrocyte-associated cells and (C) NeuN⁺ neuronal populations, showing reduced cell recovery in CDD-fed mice. (D) Schematic of hippocampal tissue collection and RT-qPCR analysis, with quantification of inflammatory and microglia-associated genes (*Ilc, Iba1, Tnfa, Cdc8, Il1b, and Ifng*). Gene expression is presented as fold change relative to GBC controls. Bars represent group means with individual biological replicates. Statistical significance: P < 0.05, P < 0.01, P < 0.001, P < 0.0001; ns, not significant.

We hypothesized that the impacts of fiber deprivation on behavior might involve changes in the hippocampus. The small size of this region relative to hemi-brain, precluded interrogating it via flow cytometry. Given this limitation, we assayed whole hippocampal gene expression via RT-qPCR **(Fig. 7D)**. We found that, relative to GBC-fed mice, increased expression of *Cd68* and *Aif1* in mice fed CDD but not CDD+INU, suggesting increased microglia. These differences were accompanied by increased expression of *Il6*, *Tnfα*, *Ifnγ*, and *Il1β*. Collectively, similar to hemibrain, these results indicate that fiber deprivation resulted in hippocampal inflammation.

## Discussion

The increasing prevalence of mental health disorders, and growing recognition that neuroinflammation contributes to such disorders, has spurred interest in identifying environmental (i.e. non-genetic) factors, including diet, that might influence neuroinflammation and/or its associated mental health states. Extensive studies of diet and gut health indicate that dietary fiber plays an important role in maintaining a healthy gut microbiota, which protects the intestine against inflammatory diseases. Viewing such findings amidst the increasing appreciation that gut health is an important determinant of brain health led us to hypothesize that lack of dietary fiber might promote phenotypes associated with mental health disorders. Our findings in part support this hypothesis. Specifically, we report that, in mice, deprivation of dietary fiber resulted in increased anxiety-like behavior, which was associated with low-grade inflammation in the brain.

Fiber deprivation was modeled by replacing the standard grain-based fiber-rich rodent chow (GBC) used in most academic vivaria with a highly refined compositionally defined low-fiber diet. However, the conclusion that this diet’s impact reflected its lack of fiber rather than one of the numerous other ways it differed from GBC was only reached after observing that specifically enriching the low-fiber diet with fiber restored behavioral patterns and neuroinflammatory readouts to that observed in GBC-fed mice. The specific fiber utilized in this regard was the fermentable fiber inulin, which was chosen based on previous studies that it, but not the fermentation-resistant fiber cellulose, restored gut health. Yet it is increasingly appreciated that dietary fibers are highly heterogenous complex molecules such that their fermentability is but one of numerous means by which they can influence gut health. Moreover, only a portion of inulin’s impacts on gut health are a consequence of its fermentation products. Thus, it will be important for future studies to determine the extent to which other fibers, including those typically consumed as supplements (e.g. psyllium) or those naturally in foods historically consumed by humans (e.g. pectin, wheat fiber), also impact behavioral aspects of mental health and, furthermore, mechanisms by which they do so.

The outwardly observable consequence of fiber deprivation was increased anxiety-like behavior as measured by the open-field test. Such increased anxiety-like behavior was associated with a broad array of indices of mild neuroinflammation including increased abundance of bone marrow derived inflammatory cells, reduced regulatory T cells, and more pro-inflammatory microglia. These changes were paralleled by reduced astrocyte function and reduced recovery of viable oligodendrocytes and neurons. We hypothesize that the changes in the immune cells drove the other observed alterations in the brain, which led to increased anxiety-like behavior. How fiber deprivation might have led to increased immune cell activation in the brain is unclear, but it might have been driven by increased abundance of gut microbiota products, or the pro-inflammatory mediators such products induce, in systemic circulation and ultimately reaching the brain. Such mediators might include metabolites like urea, which is reported to have neurological impacts, and which was reduced in serum upon enrichment of the low-fiber diet with inulin but could also include innate immune activators such as lipopolysaccharide (LPS) [22,23]. Levels of such mediators in serum could be elevated due to increased production and/or increased gut permeability. The latter possibility essentially holds that fiber deprivation-induced neuroinflammation was a consequence of its degradation of gut health. Concomitantly it predicts that fibers such as cellulose, which do not rescue gut health when added to low-fiber diets, would not rescue brain health. In contrast, adding either psyllium or wheat fiber to low-fiber diets protect against gut inflammation suggesting they might also provide neuroprotective benefits. Impacts of inulin, psyllium and wheat fiber are all mediated by gut microbiota. However, impacts of these fibers on microbiota differ dramatically as do the mechanisms by which they promote gut health suggesting that each fiber may impact the serum metabolome, and consequently, the brain in a unique manner. Thus, further studies to examine the extent to which these fibers impact the brain are warranted.

In conclusion we report that, in mice, dietary fiber deprivation results in neuroinflammation and increased anxiety-like behavior. An important caveat to these findings is that our study only utilized male mice and thus, relevance to female mice, let alone humans is unknown. Nonetheless, should our findings prove applicable to humans, it would indicate that dietary fiber is not only important for gut health but that insufficient fiber consumption can negatively impact brain health. Our results support the concept of consuming prebiotic fiber supplements as a means of promoting brain health. However, we caution that, at least in mice, inulin-enriched diets can have detrimental consequences not observed for diets naturally rich in fiber. Hence, at least until extensive human studies are performed, we suggest consumption of foods naturally rich in fiber as the best means of promoting gut and perhaps brain health.

